# Multi-scale machine learning model predicts muscle and functional disease progression in FSHD

**DOI:** 10.1101/2025.02.19.639046

**Authors:** Silvia S. Blemker, Lara Riem, Olivia DuCharme, Megan Pinette, Kathryn Eve Costanzo, Emma Weatherley, Jeff Statland, Stephen J. Tapscott, Leo H. Wang, Dennis W.W. Shaw, The Move+ Investigator Team, Xing Song, Doris Leung, Seth Friedman

**Author notes:** Corresponding Author: Silvia S. Blemker, Ph.D. E-mail Address Address: 110 Old Preston Ave., Charlottesville, VA, USA 22902.

## Abstract

Facioscapulohumeral muscular dystrophy (FSHD) is a genetic neuromuscular disorder characterized by progressive muscle degeneration with substantial variability in severity and progression patterns. FSHD is a highly heterogeneous disease; however, current clinical metrics used tracking disease progression lack sensitivity for personalized assessment, which greatly limits the design and execution of clinical trials. This study introduces a multi-scale machine learning framework leveraging whole-body magnetic resonance imaging (MRI) and clinical data to predict regional, muscle, joint, and functional progression in FSHD. The goal this work is to create a ‘digital twin’ of individual FSHD patients that can be leveraged in clinical trials.

Using a combined dataset of over 100 patients from seven studies, MRI-derived metrics—including fat fraction, lean muscle volume, and fat spatial heterogeneity at baseline—were integrated with clinical and functional measures. A three-stage random forest model was developed to predict annualized changes in muscle composition and a functional outcome (timed up-and-go (TUG)). All model stages revealed strong predictive performance in separate holdout datasets. After training, the models predicted fat fraction change with a root mean square error (RMSE) of 2.16% and lean volume change with a RMSE of 8.1ml in a holdout testing dataset. Feature analysis revealed that metrics fat heterogeneity within muscle predicts muscle-level progression. The stage 3 model that combined functional muscle groups and predicted change in TUG with a RMSE of 0.6 seconds, in the holdout testing dataset. This study demonstrates the machine learning models incorporating individual muscle and performance data can effectively predict MRI disease progression and functional performance of complex tasks, addressing the heterogeneity and nonlinearity inherent in FSHD. Further studies incorporating larger longitudinal cohorts as well as comprehensive clinical and functional measures will allow for expanding and refining this model. As many neuromuscular diseases are characterized by varability and heterogeneity similar to FSHD, such approaches have broad applicability.

## INTRODUCTION

Facioscapulohumeral muscular dystrophy (FSHD) is a genetic neuromuscular disorder characterized by progressive muscle weakness and wasting, affecting approximately 1 in 7,500 individuals^1^. The genetic basis for FSHD—mutations leading to aberrant expression of the protein DUX4, which is toxic to muscles—is well established. Disease progression typically occurs over years in adult-onset FSHD and can affect skeletal muscles across the entire body ^2–10^. However, there are no approved treatments for FSHD, and no effective approaches exist to mitigate the progressive weakness and associated loss of key functions such as ambulation, reaching, and fascial expression.

Current clinical metrics for tracking FSHD progression primarily rely on functional assessments, including manual muscle testing, timed movement tests (e.g., timed up-and-go (TUG)), computer-vision-analyzed movements (e.g., reachable workspace^11^), and patient-reported outcomes. While these measures can provide valuable insights into function over longer periods, they have significant limitations in shorter-term settings, such as clinical trials lasting for shorter periods of time, such as a year or less. Functional measures often exhibit inherent variability from test to test and day to day, reducing their reliability for detecting meaningful changes over short durations. Furthermore, the heterogeneity of FSHD means that the most relevant functional task for tracking disease progression likely differs across patients. For instance, only a subset of participants in a clinical trial may experience measurable declines in shoulder girdle strength, impacting their RWS performance, particularly in the absence of treatment. The limitations of such measures were recently highlighted in the Fulcrum Therapeutics Phase III trial (NCT05397470), where both placebo and treatment groups showed improvements in RWS, ultimately preventing the detection of meaningful treatment effects. The reasons for this placebo effect remain unclear but could involve motor learning effects or placebo responses. Similar limitations extend to other functional metrics, such as TUG and the 6-minute walk test, which may not capture progression uniformly across all patients.

To address the limitations of global clinical metrics, magnetic resonance imaging (MRI) has emerged as the gold standard for quantifying muscle-level involvement in FSHD. MRI enables a detailed assessment of muscle composition, particularly the amount of fat infiltration, which serves as a reliable metric of disease expression. Despite the relative ease of acquiring whole-body imaging data, current analytic methods for segmenting and quantifying features at the individual muscle level remain limited. Consequently, much of the existing literature in adults ^3–5,8,10,12–24^ and children ^25,26^ has relied on qualitative ratings or quantification confined to single slices or central muscle sub-regions. Recent advances have introduced semi-automated ^21^ and fully automated methods ^22,24^ to quantify select individual muscles or combined muscle groups, leading to updated conclusions about muscle involvement and fat progression patterns. Key revisions include: (1) while FSHD was historically thought to primarily affect specific muscles (e.g., scapular fixation muscles, hamstrings, tibialis anterior, and gastrocnemius), it is now evident that all muscles can be affected across the lifespan; (2) the pattern of muscle involvement and progression is highly variable across patients and muscle groups; and (3) within individual muscles, disease progression is heterogeneous, with varying degrees of fat infiltration and diverse patterns of structural degradation.

As trials have progressed—such as the recently completed trial (NCT05397470) and ongoing studies (NCT05747924 and NCT05548556)—the common approach has been to monitor a subset of individual muscles or combined muscle groups. This involves grouping muscles with a prespecified range of fat infiltration, deemed “at risk,” into a single category and using this aggregated number as a treatment response metric ^22^. While this streamlined strategy aligns with the single-biomarker approach used in DMD ^27^, it has several notable limitations: (1) it omits muscles above and below the specified threshold that may also be changing; (2) it disregards baseline muscle status and identity, which are known to influence change rates; (3) it overlooks fat distribution patterns, with confluence of fat being a characteristic feature of the disease; and (4) it averages data in a way that pulls results toward a central trend, masking individual variability and reducing the sensitivity needed to detect subtle but clinically significant changes. Collectively, these limitations diminish the ability of MRI to capture nuanced disease progression and weaken its utility in correlating muscle changes with functional outcomes.

While a whole-body MRI analysis provides a more in-depth, personalized description of FSHD disease state, the vast amount of data generated also introduce significant analytical challenges. Furthermore, muscle MRI measurements do not provide the full picture of an individual’s disease state: other data types, such as clinical assessments and disease descriptors (both descriptive modulators like D4Z4 allele length and functional task performance), can also be incorporated to provide an integrated patient assessment tool. These challenges point to the need for updated methods beyond traditional parametric statistics to interpret and analyze the data effectively. Machine learning techniques, particularly ensemble methods, have shown considerable promise in addressing the complexities of heterogeneous disease in order to predict patient-specific disease progression. Indeed, machine learning has been applied in the context of clinical trials for other diseases through synthetic control arms ^28,29^ and AI-driven digital twins of patients ^30^. These examples highlight the increasing role of AI-based models in addressing disease heterogeneity, which remains a major challenge in FSHD trials.

The primary objective of this study is to develop a machine learning model that incorporates detailed MRI measures along with clinical data and use the model to predict and advance our understanding of FSHD disease progression. Beyond individual outcome prediction, our goal is to create an FSHD patient digital twin—a model that simulates the natural course of disease progression in untreated patients.

## METHODS

### Data Collection

We leveraged multiple FSHD retrospective datasets, combined into a “data lake.” The datasets included MRI scans collected from >100 patients with FSHD from seven different studies: Wellstone ^31^ cohort (n = 34, P50 AR065139, J. Chamberlain PI, S. Tapscott co-PI); Kennedy Krieger Institute (KKI) cohort (n = 30)(1K23NS091379, D. Leung PI), a subset of the Fulcrum Phase II DUX4 placebo cohort (n = 20, NCT05397470), FSHD Global Research Foundation Registry cohort (n = 28), and MOVE+ patients (n = 24) (Statland PI, funded by Avidity Biosciences, FSHD Canada). All experimental protocols used to collect the data were in accordance with relevant guidelines/regulations and were approved by local or central IRBs. Subjects provided informed consent for data collection and aggregation, consistent with the local or central IRBs at all the same institutions. The scans varied in coverage and acquisition method (Table 1).

**Table 1:** Data set information.

| **Site** | **Region of Coverage**<br>*Distribution of scans* | **# of Subjects**<br>*Sex Distribution* | **Timepoint Distribution**<br>*# Scans per Timepoint* | **Height (cm)**<br>*Mean +/- Stdev (Range)* | **Weight (kg)**<br>*Mean +/- Stdev (Range)* |
| --- | --- | --- | --- | --- | --- |
| **BILAT** | LE<br>(96) | 34<br>*M - 15, F - 19* | Baseline: 34<br>52 Weeks: 30<br>104 Weeks: 27<br>156 Weeks: 5 | 175.11 +/- 11.34<br>(152.0 - 201.0) | 81.08 +/- 13.63<br>(53.4 - 111.0) |
| **KKI** | LE/CO<br>(69/73) | 30<br>*M - 12, F - 18* | Baseline: 30<br>12 Weeks: 27<br>36 Weeks: 26<br>60 Weeks: 30<br>84 Weeks: 29 | 173.13 +/- 11.29<br>(154.0 - 196.0) | 81.94 +/- 19.18<br>(47.8 - 118.9) |
| **MOVEPlus** | LE/FB<br>(19/13) | 24<br>*M - 20, F - 4* | Baseline: 24<br>52 Weeks: 8 | 177.63 +/- 10.42<br>(157.0 - 193.0) | 82.81 +/- 16.22<br>(55.7 - 122.3) |
| **Fulcrum** | LE/FB<br>(21/39) | 20<br>*M - 15, F - 5* | Baseline: 20<br>48 Weeks: 20<br>96 Weeks: 20 | 178.05 +/- 11.21<br>(152.4 - 199.0) | 86.0 +/- 15.44<br>(54.9 - 119.0) |
| **FSHD Global** | FB<br>(29) | 29<br>*M - 17, F - 12* | Baseline: 29 | 175.25 +/- 10.78<br>(155.0 - 200.0) | 78.18 +/- 15.73<br>(55.0 - 110.0) |

### MRI Segmentation

We utilized an AI-based approach similar to our previously published algorithm and described methods ^32^ to segment the boundaries of up to 118 muscles (depending on coverage) across the whole body from the Dixon water images. As per our prior work, trained segmentation engineers manually reviewed and edited for accuracy (process we call “vetting”) utilizing the manual editing tools (i.e. paint, draw, erase) in 3D Slicer (v4.11). This process required between one and three hours, depending on the level of fatty replacement (with higher fatty replacement muscles requiring more user interaction time for contour verification).

### Muscle distribution metrics

Since disease progression is often heterogeneous within each muscle in FSHD, the Stage 1 model focused on regional level metrics. First, we computed muscle and fat quantities within each axial slice of each muscle and displayed those metrics as a function of length along the muscle (as described in ^24^). Slice-level metrics have the advantage of greater sensitivity than overall metrics, given that fat is often regionally localized and changes occur within that limited region. This step yieleded individual axial slice measures of muscle boundary cross-sectional area (CSA), lean muscle CSA, and area fat fraction (%), all of which are relatively standard metrics used in the field. Additionally, we calculated three metrics of heterogeneity and distribution of fat within each slice: fat variation, kurtosis, skewness, and Moran’s index. Fat variation (in units of %) for eachregion was calculated as the standard deviation of fat fraction across all pixels in the boundary volume. Kurtosis and Skewness (in units of %) for each region were calculated as the kurtosis and skewness across all pixels in the boundary cross-section. Moran’s index (normalized) for each region assesses whether nearby or neighboring areas are more similar (or dissimilar) than what would be expected under random spatial distribution; our implementation was based on Santago et al.^33^ The measures were then expressed as a function of longitudinal distance slice-by-slice moving inferior (distal) to superior (proximal). Muscle characteristics were then expressed as a function of the percentage of the muscle length from 0% (inferior end of the muscle) to 100% (superior end of the muscle) and were sampled at 10% increments.

### Muscle-level metrics derived from MRI data

Our prior work ^32^ demonstrated the wealth of information that can be gleaned from analysis of segmentations of muscles in patients with FSHD. These findings led us to incorporate multiple regional-level metrics into the multi-scale model. Each muscle analysis was performed on each image, expressed along the superior-inferior length of the muscle, as previously described ^32^; curves were sampled to produce metrics within 8 regions of each muscle (Table 2). Specific values were calculate based on the following definitions. Fat fraction (FF: in units of %) for each muscle was determined from the combination of fat and water images. Lean muscle volume (LMV: in units of ml) was determined by subtracting the fat volume from the boundary volume (BV): LMV = BV – FF*BV.

**Table 2:** Model parameters.

| **Stage 1. Regional-Level Model** |  |
| --- | --- |
| **Parameter** | **Description** |
| *Region Fat Fraction* | Fat fraction within a region |
| *Region CSA* | Region CSA |
| *Relative CSA* | Ratio of region CSA to average muscle CSA |
| *Difference From FFneighbors* | Difference between region FF and neighboring regions FF |
| *FF Range Region* | Max regional FF - Minimum regional FF |
| *Region FF Variation* | Standard deviation of FF in region |
| *Region FF Skewness* | Skewness of FF in region |
| *Region FF Kurtosis* | Kurtosis of FF in region |
| *Region FF Moran* | Moran's Index of FF in region |
| **Stage 2. Muscle-Level Model** |  |
| **Parameter** | **Description** |
| *Region Prediction* | Output of stage 1 prediction |
| *Fat Fraction Z score* | Z score of fat fraction relative to FSHD population at baseline |
| *Fat Fraction* | Fat fraction at baseline |
| *Higher ranking muscles score* | Average of Fat fraction z scores in muscles with higher ranking |
| *Lower ranking muscles score* | Average of Fat fraction z scores in muscles with lower ranking |
| *LMV to bone Z score* | Z score of lean muscle volume to bone CSA, compared to FSHD population |
| *Femur CSA* | Cross sectional area of femur |
| *Fat fraction range across regions* | Max fat fraction across sub regions – min fat fraction across sub regions |
| *Femur Fat Fraction* | Fat fraction within femur |
| *D4Z4 repeat length* | D4Z4 repeat length (integer value) |
| *Bone to Height Z-score* | Z score of bone to height |
| *TUG time* | Time to up and go |
| *Muscle* | Muscle name (classifier) |
| **Stage 3. Functional-Level Model** |  |
| **Parameter** | **Description** |
| *TUG at Baseline* | Timed up-and-go at baseline |
| *Knee extensors normalized change in LMV from baseline* | Output from combined prediction of Stage 2 LMV model, normalized to height*mass |
| *Hip abductors normalized change in LMV from baseline* | Output from combined prediction of Stage 2 LMV model, normalized to height*mass |
| *Hip extensors normalized change in LMV from baseline* | Output from combined prediction of Stage 2 LMV model, normalized to height*mass |
| *Hip flexors normalized change in LMV from baseline* | Output from combined prediction of Stage 2 LMV model, normalized to height*mass |
| *Trunk extensors normalized change in LMV from baseline* | Output from combined prediction of Stage 2 LMV model, normalized to height*mass |

### Bone metrics

The volume, average cross-sectional area, and fat fraction of each femur bone was determined. The CSA and volume were used to standardize muscle volume metrics and fat fraction was incorporated because prior work has demonstrated a correlation between trabecular fat fraction and bone mineral density measured by quantitative computed tomography^34^.

### Analysis of baseline MRI muscle data for model feature development

The baseline muscle fat fraction and lean muscle volume data were analyzed to examine trends across the FSHD population and to establish normative ranges for metrics within the FSHD population. Analysis included: fat fraction for each muscle, fat variation for each muscle, and lean volume (adjusted for body size) for each muscle. In addition to standard descriptive statistics, heatmap visualizations of muscle involvement, with elements organized by the average fat fraction across muscles (within each patient) and TUG times were used to characterize the relationships between pattern of muscle involvement and overall functional and global fat replacement. The relationships between total volume and the product of femur cross-sectional area and lean muscle volume across patients with FSHD were determined, to size adjusted value.

Based on these analyses, two metrics were generated. First was an estimation of the degree to which each muscle is following a global pattern of progression across all patients. For each muscle, the composite fat fraction of the subset of muscles that were determined to be more frequently involved across patients was computed for that particular patient. If this value is low then this muscle is less likely to change, whereas, if the value is high, then this muscle is more likely to change. Second was an estimation of the likelihood of change of a muscle based on a distribution analysis of the baseline fat fraction values for that particular muscle.

### Clinical and functional data

The D4Z4 repeat length and timed up-and-go (TUG) was included in the baseline description of each patient. For a subset of patients, TUG measurements were also available at follow up time points. All subjects that had longitudinal MRI measures and thus were simulated with the disease progression model were confirmed FSHD Type 1. Unfortunately, the metrics of disease severity were not the same across all cohorts and thus were not used in the model. Individual patient age and sex were not included in the model because: 1) preliminary analysis indicated that they were not features of high significance, and 2) excluding these features allowed for maximal use of the data because age and sex were not known for all patients.

### Three-Stage, Multi-scale Model Development

The random forest algorithm was chosen for its ability to handle complex, non-linear relationships and its resistance to overfitting. Prior research has established the suitability of this approach for predicting disease progression^35,36^. To incorporate both regional and total muscle metrics, we developed a multi-stage random forest training and validation approach (Fig. 1). The first two models are designed to predict individual muscle progression over time, with the third model designed to incorporate the output from the first two model stages to predict change in TUG performance. The first stage, one unique random forest model, included training on a subset of the data to predict the changes in fat fraction and lean cross-sectional area at the regional level. This allows the model to capture how the local tissue structure influences progression of fat infiltration and atrophy in a specific region. This model was trained on 10% of the full data set, which included 6330 muscle regions. The data used for the stage 1 model training was not included in any other training or testing datasets. The second stage, a two parallel-running random forest model, included training another subset of the data (that was not included in the prior training datasets) to predict the whole-muscle level changes in fat fraction and lean muscle volume. Each of these samples were first run through the stage 1 model to predict regionalized changes; then the cross-sectional area weighted average of the regional fat fraction changes was calculated and inputted into the stage 2 model. This model was trained on 70% of the full data set, which included 4513 muscles, and the model was tested on 833 muscles. The outputs of the stage two models were combined to calculate functional group changes, by calculating volume-weighted average of the individual fat fractions at the joint level, and summing the lean muscle volumes, normalized by the product of height and mass. Finally, these right and left group-level predicted changes averaged and incorporated into the stage 3 random forest model to predict change in TUG measurements. The stage 3 model was trained with 26 data points and tested with 4 data points (only a subset of patients included had both coverage of major lower limb muscle groups as well as multiple measurement of TUG).

**Figure 1:**
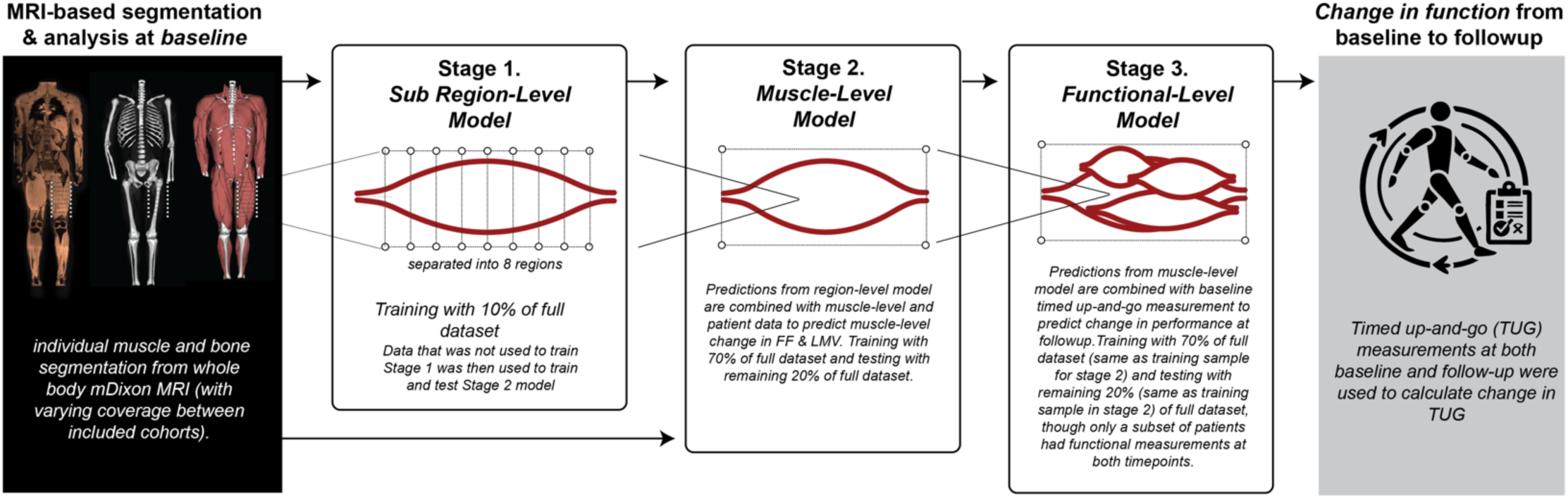
Illustration of the multi-scale disease progression model. Three separate stages which examined specific scales of muscle involvement were developed. Outputs from each stage were used to determine inputs to the next stage.

For each stage, a random forest model consisting of around 500 trees was trained on the training dataset. Models were implemented in MATLAB (Mathworks, Natick, MA, USA), using the combination of templateTree and fitrensemble function. Grid search was conducted to optimize key hyperparameters such as the number of trees, maximum depth, and minimum samples per split. The out-of-bag method was used to prevent overfitting. Shapley Additive exPlanations (SHAP) analysis^37,38^ was used to identify the key variables contributing to predicted outcome.

### Model Evaluation

Each model was evaluated by assessing the ability to predict annualized change in fat fraction for each region (for Stage 1), muscle-level fat fraction or lean muscle volume (for Stage 2) for both the training and testing data samples. The Stage 3 model was evaluated by assessing the ability to predict change in timed up-and-go. The performance accuracy metrics included: mean error (ME), root-mean-square error (RMSE), as well as Pearson’s correlation coefficient between predicted and actual fat fraction or lean muscle volume change (r) and its p-value. Feature importance scores were extracted from the random forest model using SHAP analysis to identify the most influential predictors of fat fraction or lean muscle volume changes.

## RESULTS

Fat fraction varied substantially across muscles and patients (Fig. 2). On average, the semimembranosus (hamstring) muscle was the most affected in the cohort while the popliteus muscle was least affected. Heatmap visualization of fat fraction across all muscles (organized average fat fraction across individuals) and patients at baseline (organized by TUG) reveal an overall pattern of muscle involvement across disease severities (Fig. 3). As TUG time increases, more muscles have fat fraction levels in the high range, consistent with the progressive nature of the disease. The order of muscles is not regional: the muscles in the ‘earlier’ group are in both upper and lower regions of the body; similarly, muscles in the ‘later’ group are muscles from all regions as well. Substantial heterogeneity across patients and muscles is most apparent in the patients in the middle range of involvement.

**Figure 2:**
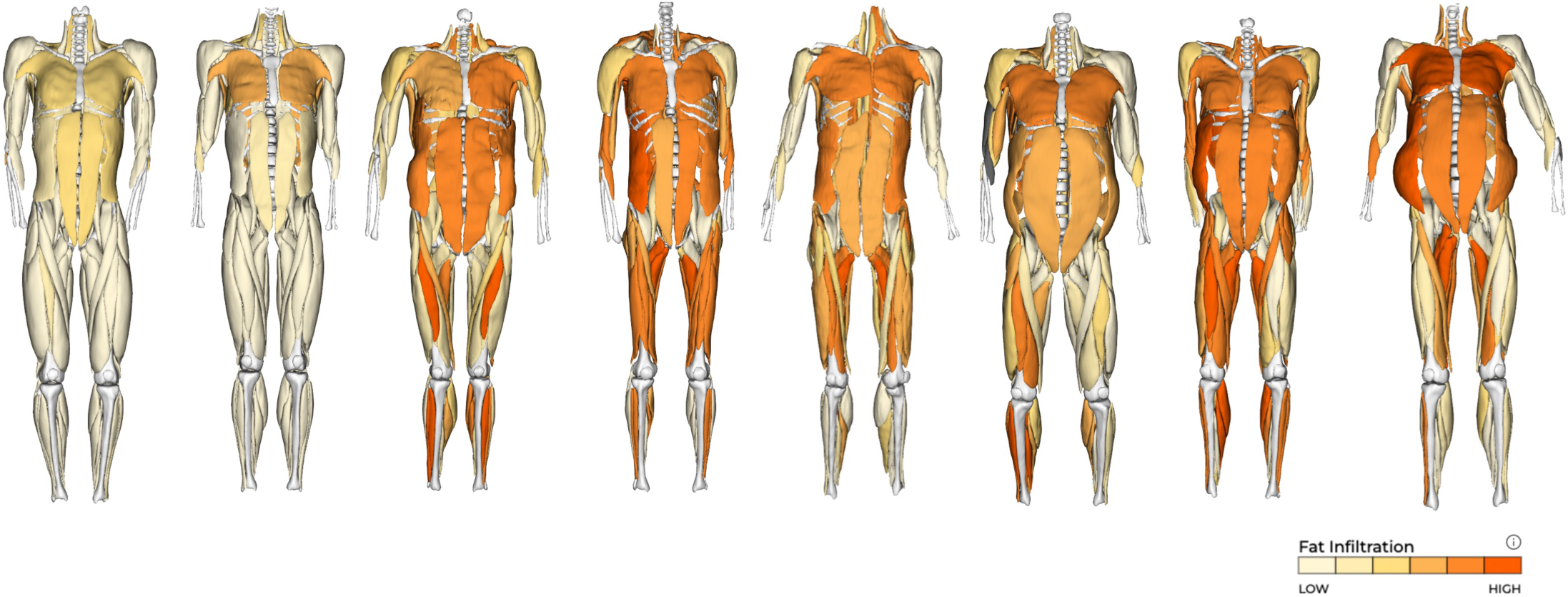
Visualization of 3D segmentations of muscles in patients with FSHD of varying disease severity. Fat fraction % is provided as a colormap on each muscle in the models to illustrate the unique pattern of involvement in each patient.

**Figure 3:**
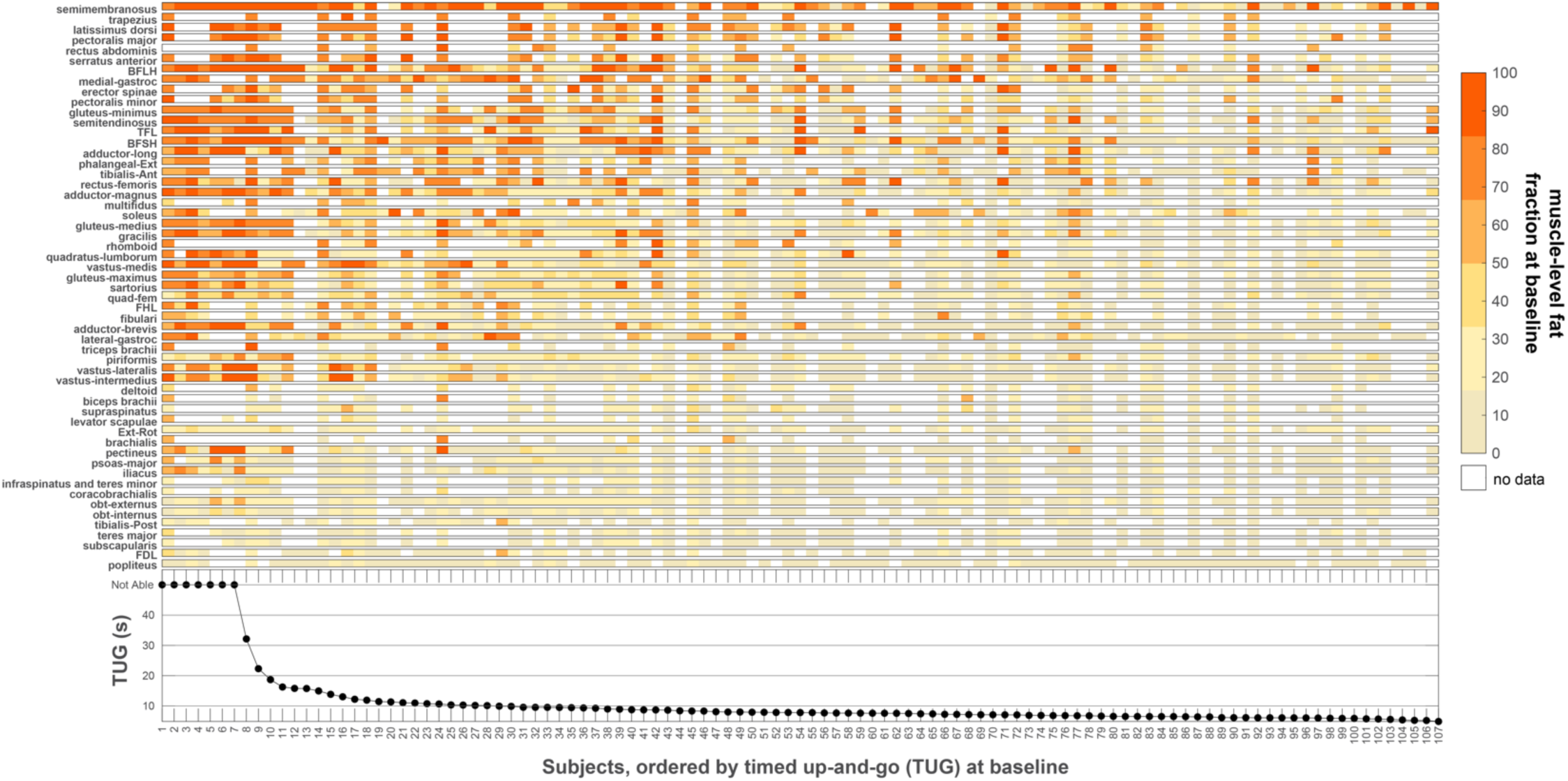
Visualization of muscle involvement at baseline in patients with FSHD, ordered by timed up-and-go measurement at baseline. The colormap is based on muscle-level fat fraction at baseline. White squares represent specific muscles that were not measured. Muscles are ordered vertically according to the average fat fraction measured across all subjects and limbs.

The Stage 1, the test prediction of regional-specific random forest model (Fig. 4) was significantly correlated with measured regional fat fraction per muscle (r = 0.43, p < 0.001), with a mean error of −0.32% and RMSE of 3.5%, within the Stage 1 testing dataset. SHAP analysis revealed the parameters: *Regional Fat Fraction Kurtosis*, *Regional Fat Fraction Variation, Difference from FF Neighbors*, and *Relative CSA* had the greatest influences on prediction of fat fraction change at the regional level.

**Figure 4:**
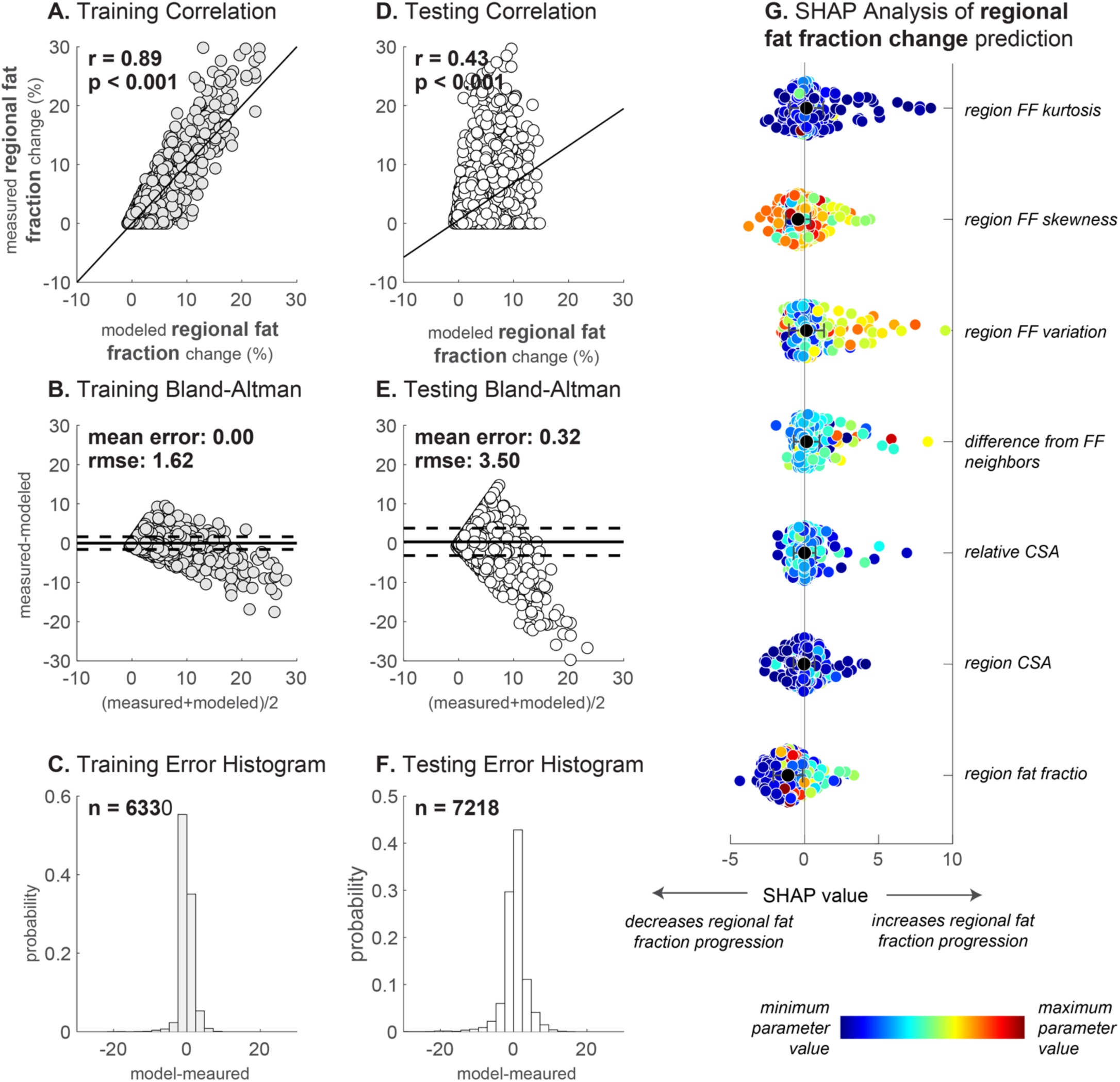
**Stage 1 Model: Regional-Level Fat Fraction Change Prediction**. Results from Stage 1 model training and testing, based on measurements at the regional level. Each muscle is separated into 10 separate regions longitudinally, and the model predicts the change in fat fraction of the region based on measurements at the regional level. The output of this stage is combined into a singular prediction (by averaging the predictions across the regions within each muscle) that serves as a variable input to the Stage 2 models.

The Stage 2, muscle-level random forest model prediction of fat fraction change (Fig. 5) was significantly correlated with measured fat fraction change in the testing dataset (r = 0.5, p < 0.001), with a mean error of −0.3% and RMSE of 2.2%. *Regional Prediction* (output from Stage 1 model), *femur cross-sectional area*, and *muscle* had the greatest influence on prediction of muscle-level fat fraction change. The Stage 2, muscle-level random forest model prediction of lean muscle volume change (Fig. 6) was significantly correlated with measured lean muscle volume change in the testing dataset (r = 0.57, p < 0.001), with a mean error of −0.8ml and RMSE of 8.3ml. *Femur cross-sectional area*, *D4Z4 repeat length*, *LMV to femur CSA z-score*, and *muscle* had the greatest influences on prediction of muscle-level lean muscle volume change. Combining the predicted individual fat fractions and lean muscle volumes to calculate functional group-level changes in fat fraction and lean muscle volume predicted the variability in functional-group level changes in the testing dataset (Fig. 7) with high accuracy. For group-level fat fraction change prediction, all RMSE values were less than 2.5%, with the exception of the trunk flexors (5.18%). For group-level lean muscle volume change prediction, the RMSE predictions varied from 3.3ml (hip external rotators) to 26.7ml (knee extensors).

**Figure 5:**
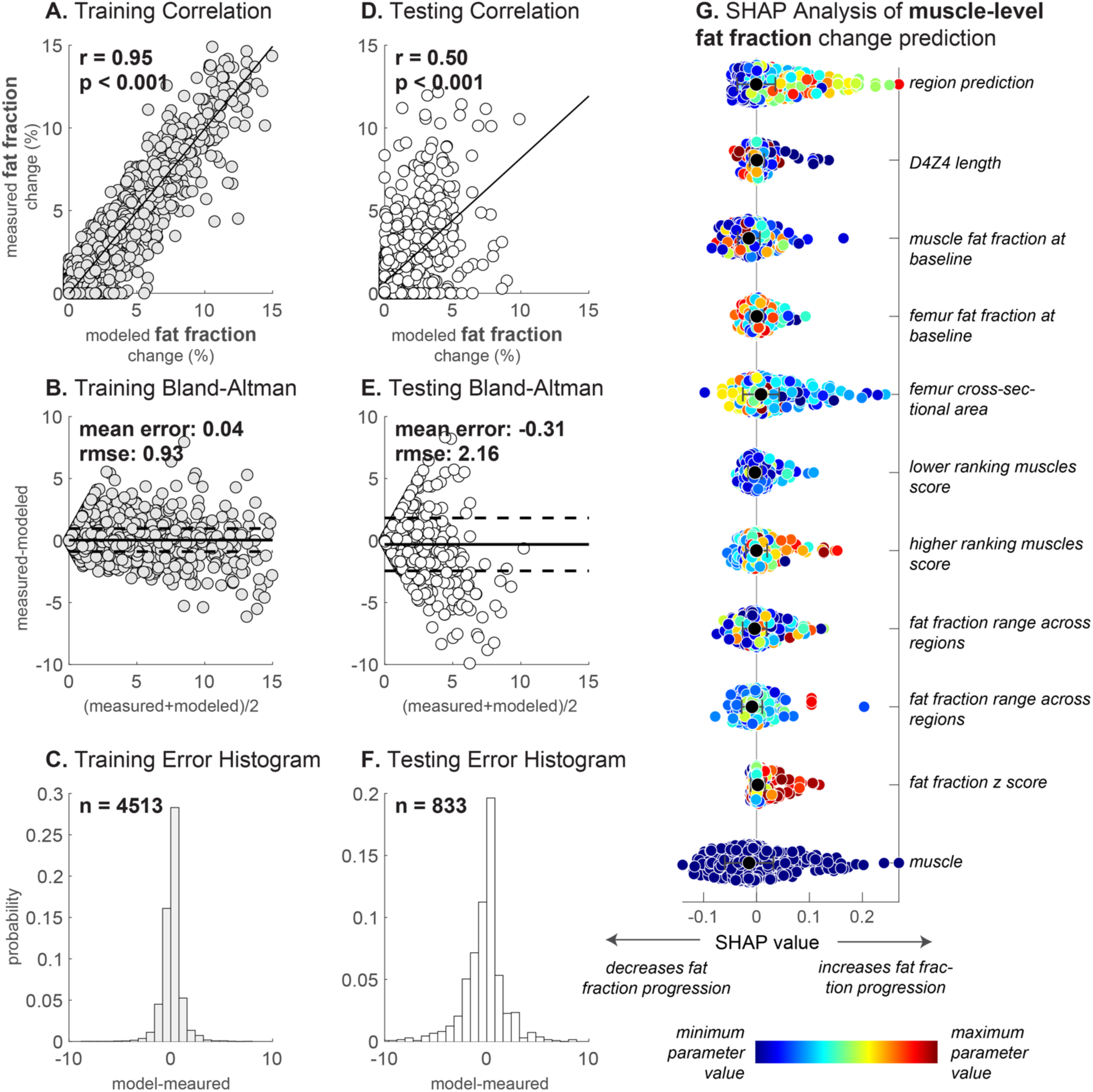
Stage 2 Model: Muscle-Level Fat Fraction Change Prediction. Results from Stage 2 model training and testing for prediction of muscle-level fat fraction changes.

**Figure 6:**
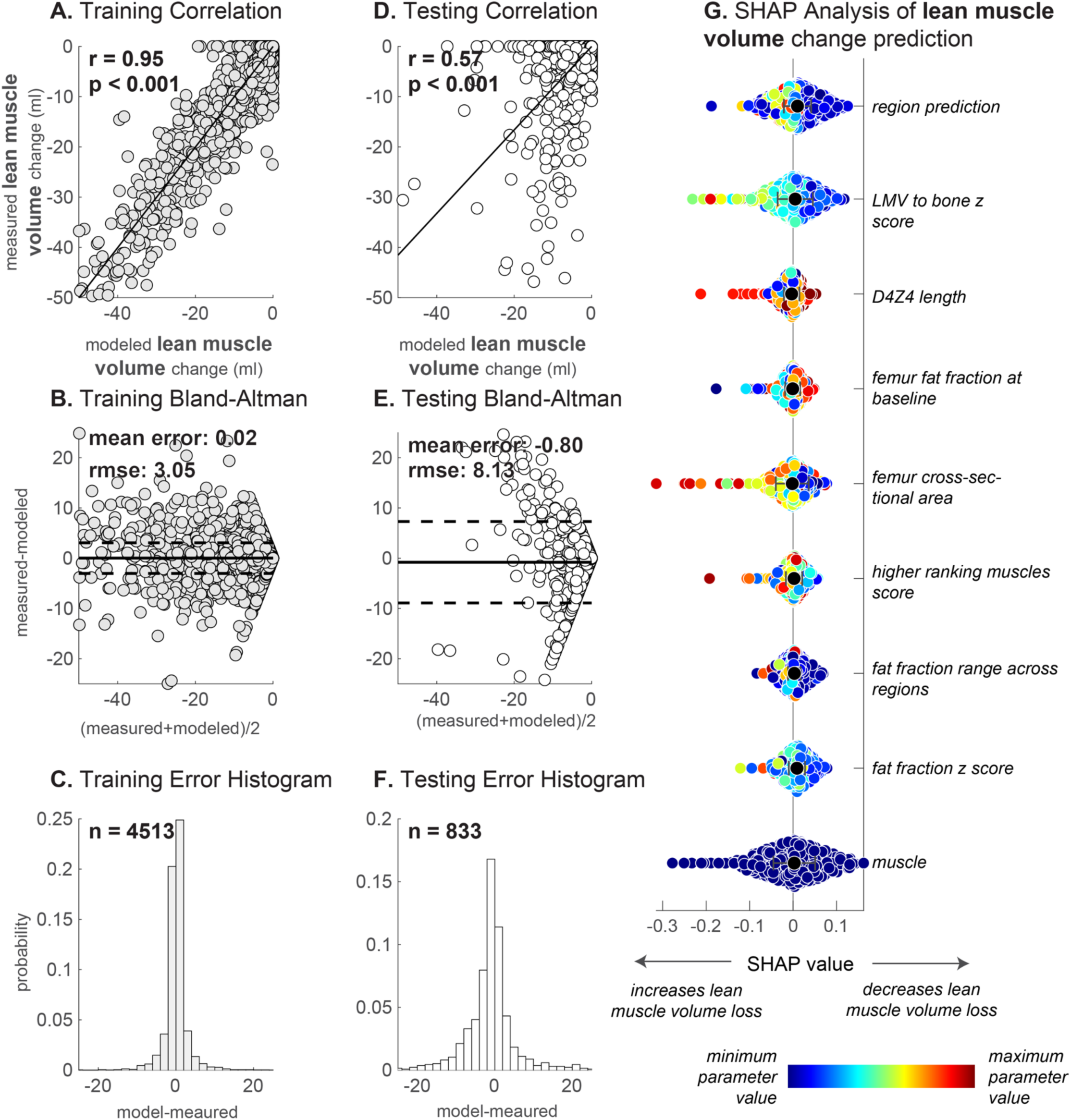
Stage 2 Model: Lean Muscle Volume Changes Prediction. Results from Stage 2 model training and testing for prediction of muscle-level lean muscle volume changes.

**Figure 7:**
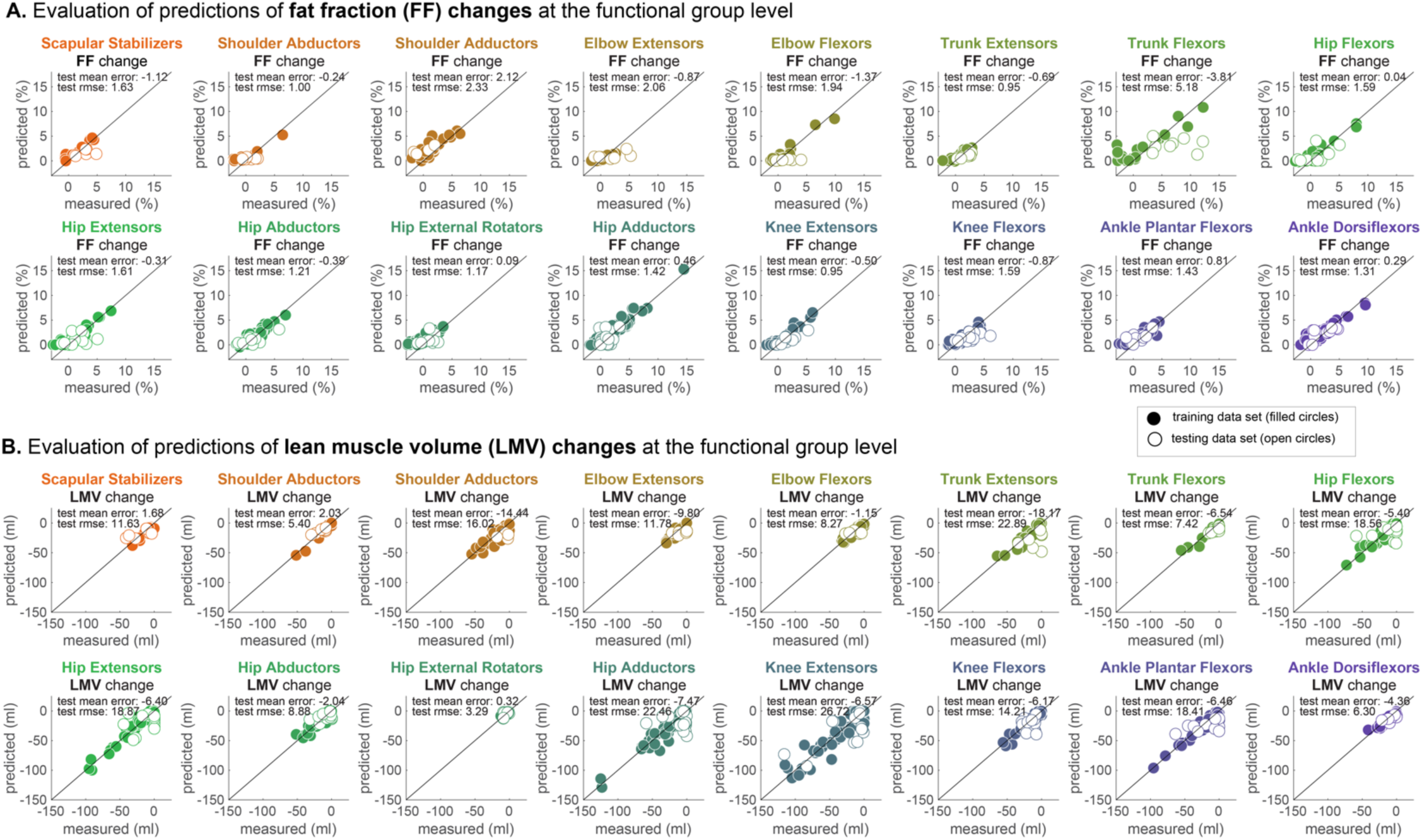
Evaluation of the output of stage 2 when determining group-level fat fraction (top two rows) and lean muscle volume (bottom two rows). Comparison between model prediction and measured values are shown for both the training datasets (solid circles) and testing datasets (open circles). Mean errors and RMSE values are reported for testing samples only. Note no errors are reported for the shoulder adductors because the dataset was limited and there were no full shoulder adductors groups in the testing dataset (which is randomly selected).

The Stage 3, functional-level random forest model prediction of change in TUG from the first to the second time point (Fig. 8) prediction had RMSE value of 0.75 s in the testing dataset. SHAP analyses revealed that the TUG at baseline had the largest influence on the predicted change in TUG, followed by the normalized change in lean muscle volume of the *Trunk Exensors*, *Hip Extensors, and Knee Extensors* muscle functional groups.

**Figure 8:**
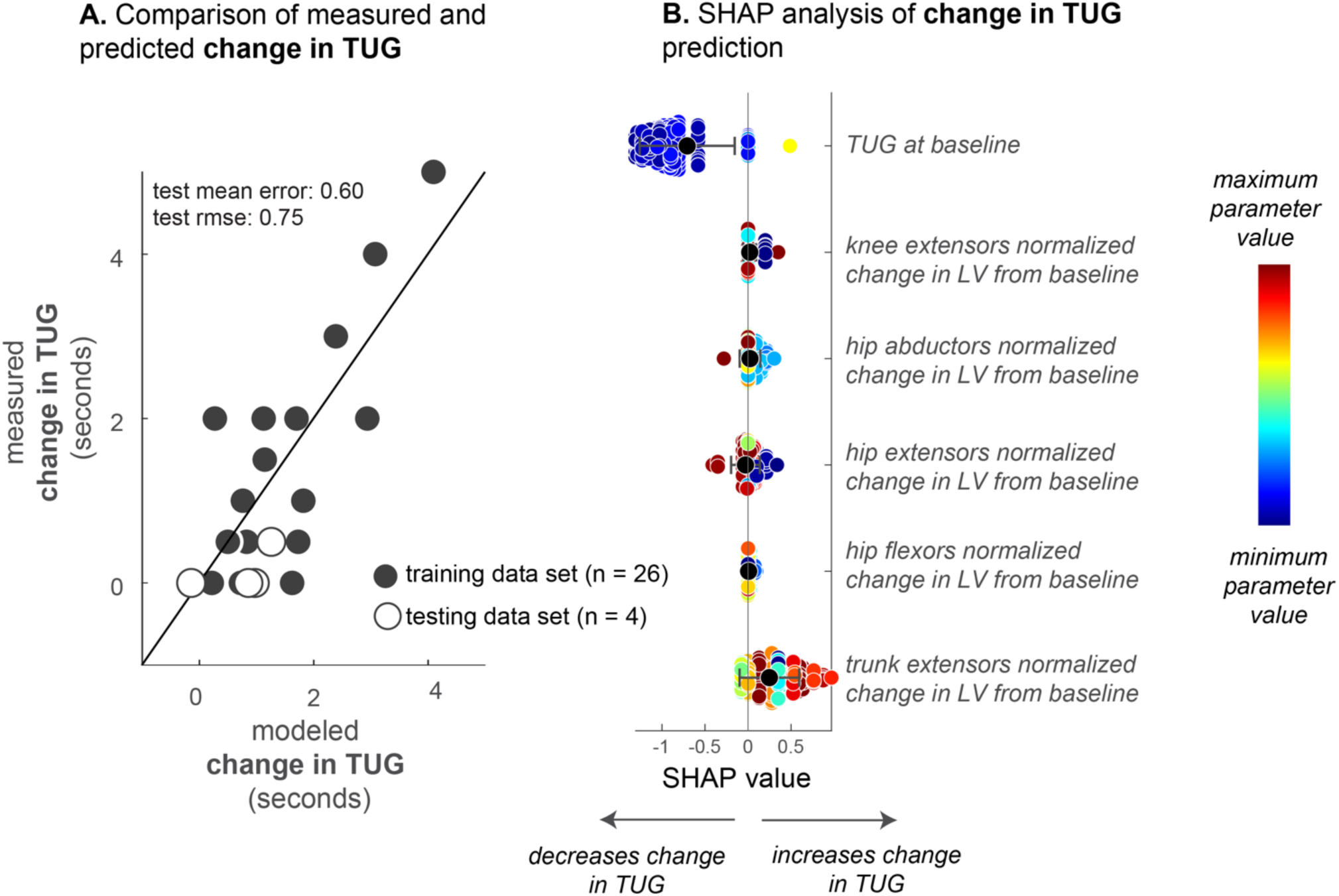
Stage 3 Model: TUG Functional Prediction. Results from Stage 3 model relating group-level normalized lean volume with timed up-and-go.,The model incorporates group-level normalized lean volume at baseline and the change in normalized lean volume from baseline the second time point to predict timed up-and-go at time point 2.

## DISCUSSION

A major challenge for drug development in neuromuscular disease is how to measure the effects of treatment in chronic progressive disorders that are marked by high variability in severity at baseline and in rates of progression. Clinical trials in FSHD currently adopt a “one-size-fits-all” approach to evaluating drug efficacy. Strategies that rely on composite metrics from MRI or performance tests risk regressing small changes to the mean, thereby reducing sensitivity in detecting meaningful differences and prolonging the required trial duration. Conversely, using a single task as a reporter is only effective if participants are pre-selected based on muscle involvement relevant to that task. Otherwise, a substantial proportion of participants—potentially unevenly distributed across placebo and treatment groups—may fail to exhibit measurable change in task performance. In this study, we present a novel ‘digital twin’ approach to predicting personalized disease progression at multiple length scales: muscle region, muscle, group, and functional levels. The developed method integrates a three-tiered machine-learning model, each component designed to capture progression at a distinct scale. This framework enables the prediction of which muscles, and to what extent, will undergo changes over time during the natural course of the disease, offering a more precise and individualized approach to monitoring FSHD progression in clinical trials. Furthemore the feature importance analyses provides insight into which factors best predict disease progression. In these models, metrics of fat fraction variability (within regions and whole muscle) were most associated with fat progression changes over one year, with bone metrics playing an important role lean muscle volume decline over one year.

Given the heterogeneity and nonlinearity of the disease, each patient has a specific set of muscles that is most likely to change at a given time. As might be expected in a heterogeneous disease, some muscle groups changed more than others in each patient, which has implications on the functional metrics that best identify disease progression in a particular patient. For example, the muscles that best predict TUG – the trunk extensors, hip extensors, and knee extensors – only changed in a subset of patients; therefore, TUG only changed in a subset of patients as well. Not surprisingly, the TUG at the baseline timepoint demonstrated a strong relationship with predicted change in TUG, as well. Taken together, these results suggest that the disease progression model provides a new approach for clinical trials that allows for patient-specific selection of functional metrics that are most likely to detect diversions from each individual’s disease progression and/or selection of qualified patients if specific functional metrics are of interest.

Analysis of the models provided insight into the parameters that influenced model prediction of disease progression. At the regional level, the initial fat fraction, the fat variation, and the kurtosis had the greatest influence on predicted fat fraction changes. These results illustrate that progression patterns are 1) muscle-specific and 2) associated with the heterogeneity of fat within the muscle. At the whole muscle level, the predicted fat fraction change from the Stage 1 model, the muscle, the D4Z4 repeat length, the associated muscle group, femur size, femur fat fraction, and metric of overall involvement influenced change in fat fraction prediction. The mechanisms underlying these observations are both intuitive and require further investigation. It stands to reason that D4Z4 length is associated with faster progression; however, the mechanisms by which bone size and composition affect disease progression representing additional complexities yet to be unraveled. Overall, these results demonstrate the complex nature of disease progression can be handled within iterative model building, allowing insights into primary predictors and individualized nature of FSHD.

The model presented in this study builds upon significant prior work in the field demonstrating general features of disease progression as determined by muscle & fat measurements on MRI ^3,6,7,10,14,15,18,21–23,39–47^. For example, a 2017 study by Andersen et al. followed 45 patients over one year, demonstrating that quantitative MRI could detect disease progression, with increased fat infiltration corresponding with declines in muscle strength and function^2^. Similarly, Fatehi et al. conducted a long-term follow-up of thigh muscles in FSHD patients, finding that quantitative MRI effectively monitored progressive fatty infiltration over time ^5^. Another common finding in the literature is that a small percentage (∼5%) of fat-affected muscles also exhibit signal elevation on short tau-inversion-recovery (STIR+) sequences ^3,7,9,47^, which can be indicative of intramuscular edema or inflammation ^3,48–50^. While this biomarker has been suggested by our group and others to foreshadow faster progression, as examples ^3,4,6,7,41^, these studies did not discriminate between muscle fat fraction at baseline, fat pattern, or muscle identity. Given the stability of the STIR+ signal when challenged with immunosuppression and steroid treatment^51^ and the priority for faster imaging exams ^52^, we believe that fat remains the optimal biomarker.

There are limitations to this study that should be acknowledged. First, while the model was developed based on likely the most complete dataset of muscle-level MRI in FSHD patients to date, more data would improve the robustness and predictive abilities of the model. Second, several parameters not included in this study would likely improve the model performance. Additional measures like methylation, further clinical assessments, and blood biomarkers represent promising possible additions to the progression model. Third, we had limited access to functional measurements at the second time point; therefore, the Stage 3 model analysis was somewhat limited and only focused on one functional measurement (TUG) and a smaller subset of subjects and muscle groups based on available longitudinal task data. Despite these factors, the model lends itself to rapid future extension, with incorporation of more functional measurements spanning lower and upper body. Lastly, the model was trained based only on adult FSHD patient data; therefore, its robustness for predicting disease progression in pediatric patients with FSHD has not been evaluated. A new training dataset, that incorporates the effects of growth assessed by a similar metric of femur-volume, would provide the opportunity for expanding the applicability of the model to pediatric populations.

This work represents a significant advancement in the use of machine learning and imaging to address the challenges of disease heterogeneity in FSHD. By integrating advanced MRI-derived muscle metrics with clinical and demographic data, the proposed multi-scale framework reveals that individual muscle progression over a year interval is predictable, given a comprensive integration of personalized data at baseline. The digital twin model serves as a powerful benchmark for assessing untreated progression, enhancing the precision and efficiency of tracking natural history changes over time. This advancement has broad potential applications in clinical trial design, including the use of personalized digital twins as surrogate placebos and leveraging the progression model’s identified features to better isolate ‘at-risk’ muscles in traditional trial designs. While incorporating additional biomarkers and expanding datasets will further refine predictive capabilities, this study establishes a strong foundation for applying machine learning and imaging technologies to neuromuscular diseases. Applying this framework has wide applicability to clinical trial design, including the use of model data to create personalized digital twins as surrogate placebos. Given variability in fat expression is a feature of many neuromuscular diseases, these methods likely have broad applicability.

## Additional Information

This work was funded by Friends of FSH Research and FSHD Global Research Foundation. Data collection of original scans was conducted as part of the NIH Wellstone (P50 AR065139, J. Chamberlain PI, S. Tapscott co-PI), K23 grants (1K23NS091379, D. Leung PI), CDC, FSHD Society, FSHD Canada, and ALSA. We thank Fulcrum Therapeutics for sharing the Phase II Redux data. Financial competing interests: Lara Riem, Olivia DuCharme, Megan Pinette, Kathryn Eve Costanzo, and Silvia Blemker are employees of Springbok Analytics. LR, EC, and OD have stock options to the company. SB is co-founder and owns stock in the company. FSHD Global is an investor in SB Analytics. JS is consultant or on advisory board for Avidity, Dyne, Fulcrum, Roche, Kate, Alnylam, Epicrispr, MiRecule, and Sanofi. SF is a consultant for Avidity, Dyne, Fulcrum, Kate, and Epicrispr.

